# Comparative electric and ultrastructural studies of cable bacteria reveal new components of conduction machinery

**DOI:** 10.1101/2023.05.24.541955

**Authors:** Leonid Digel, Mads L. Justesen, Robin Bonné, Nico Fransaert, Koen Wouters, Pia B. Jensen, Lea E. Plum-Jensen, Ian P. G. Marshall, Louison Nicolas-Asselineau, Taner Drace, Andreas Bøggild, John L. Hansen, Andreas Schramm, Espen D. Bøjesen, Lars P. Nielsen, Jean V. Manca, Thomas Boesen

## Abstract

Cable bacteria encompass at least two genera, and they are known to vary greatly in habitat preferences and filament thickness. We systematically investigated variations and similarities in cellular structures and electrical properties of different cable bacteria strains. Using SEM, TEM, STEM-EDX and ToF-SIMS, we characterized shared features of cable bacteria, such as inner and outer membranes, surface layer and cell junction architecture, as well as strain specific features, like the number and size of periplasmic conductive fibers (PCFs). Our data indicates that the PCFs are organized as loose stranded rope-like structures. With spatially resolved elemental analysis we detected nickel-containing co-factors within the PCF of cable bacteria strains in both genera suggesting a conserved conduction mechanism. Electrical conductivity of different cable bacteria strains showed a range of values covering three orders of magnitude indicating an unknown metabolic adaptation. Using cryogenic electron tomography we discovered multiple polar chemosensory arrays, abundant cytoplasmic inner membrane-attached vesicles (IMVs), polysomes and inner membrane invaginations that shed light on cable bacteria metabolism including complex motility control mechanisms, localized protein synthesis, and membrane remodeling. We propose that the IMVs discovered in this work are novel metabolic hubs closely connected to the unique conductive fiber structure of cable bacteria.

## Introduction

Cable bacteria form a diverse group of filamentous microorganisms with an ability to conduct electrons over centimeter distances (Pfeffer *et al*. 2012; Sereika *et al*. 2023). They are present globally in aquatic sediments and all cable bacteria are currently classified into two candidate genera, members of which are commonly found in freshwater or saltwater habitats (Dam *et al*. 2021; Sereika *et al*. 2023). Cable bacteria couple spatially separated sulfide oxidation with oxygen reduction via periplasmic conductive fibers (PCF) (Pfeffer *et al*. 2012; Cornelissen *et al*. 2018; Thiruvallur Eachambadi *et al*. 2020). Thousands of cells can be wired together to form centimeter-long filaments with exceptional conductivity exceeding 10 S/cm, comparable to organic semi-conductors (Le *et al*. 2017; Meysman *et al*. 2019; Bonné *et al*. 2020, 2022). The PCFs are located inside conspicuous parallel longitudinal ridges on the bacterial surface that span across cell junctions (Pfeffer *et al*. 2012). To date, little is known about the structure of the PCF beyond their approximate dimensions and localization (Cornelissen *et al*. 2018). The ultrastructural details of the whole conductive machinery have not been elucidated, and only two publications have reported on the unique elemental composition of a single marine cable bacteria species originating from Rattekaai salt marsh (Boschker *et al*. 2021; Thiruvallur Eachambadi *et al*. 2021). Based on a low-resolution mass spectrometry approach, the authors proposed a structural model, according to which cable bacteria conduct electrons through protein-based fibers with a nickel-sulfur ligated protein in their core. The conductive core was suggested to be covered by another protein layer forming an insulating surface of the PCFs. Boschker and co-workers found that treatment of cable bacteria with the surfactant sodium dodecyl sulfate (SDS) would remove the inner and outer membranes and the cytoplasm but leave the entire PCF network bound by a polysaccharide-rich base layer intact (cable bacteria skeletons). Notably, the results indicated that nickel is involved in the long-range electron transport, which is remarkable given that biological electron transport usually involves iron and copper metalloproteins (Liu *et al*. 2014).

It was previously shown that cable bacteria, despite having different sizes, have similarly sized PCFs (Cornelissen *et al*. 2018). Based on this, we hypothesized that the structure and composition as well as the electrical conductivity of the PCFs should be similar among different genera and species of cable bacteria. Electrical conductivity measurements revealed similarity of the current-voltage profiles between cable bacteria strains, however, a universal conductivity level was not observed. Notably, two of the marine cable bacteria species showed substantially lower conductivity in comparison to other species tested, indicating that PCF conductivity could be adapted to the metabolic needs of individual species in different habitats. We employed Scanning Transmission Electron Microscopy with Energy-Dispersive X-ray spectroscopy (STEM-EDX) and Time-of-Flight Secondary Ion Mass Spectrometry (ToF-SIMS) analyses to study the elemental composition of *Ca*. Electronema aureum GS (GS) and *Ca*. Electrothrix communis RB (RB) cable bacteria and showed that indeed nickel is present in the PCF of both major cable bacteria genera. In addition, high resolution cryogenic Electron Tomography (cryo-ET) combined with Transmission Electron Microscopy (TEM) and Scanning Electron Microscopy (SEM) on intact cable bacteria cells and extracted PCF allowed us to compare morphology of different strains. We accurately measured the dimensions of the PCF and investigated the details of the membrane and the cell junction lamellae ultrastructure. Extracellular filaments resembling type IV pili were observed protruding from cable bacteria using cryo-ET. Based on their position at cell-cell junctions we suggest that they play a role in gliding motility, which could be coupled to signals from adjacent chemosensory arrays. Finally, our detailed ultrastructural studies revealed highly novel and intriguing IMVs localized close to the PCFs. We hypothesize that important metabolic machinery could be associated with the IMVs forming a unique metabolic environment that is directly connected to the PCF structure.

## Results

### Variability in electrical conductivity of different strains

Cable bacteria in this study belonged to two main phylogenetic branches residing in freshwater and marine habitats respectively. The freshwater cable bacteria *Ca*. Electronema aureum GS (GS) and the marine cable bacteria *Ca*. Electrothrix communis RB (RB) were established as single strain enrichment cultures (Thorup *et al*. 2021 and this work) (Figure 1A). In addition, cable bacteria from Rattekaai salt marsh (Rat, (Meysman *et al*. 2019)) and from Hou beach (Hou and Hou1) were taken from environmental samples with mixed microbial communities. In total, we measured and compared electrical conductivity of five different cable bacteria samples to assess the variability of their conductivity and the potential role of this diversity on their metabolism. In order to understand the uniqueness of cable bacteria conduction, we measured the conductivity of 12 filamentous bacteria pure cultures and 3 from environmental samples using interdigitated gold electrodes. The conductivity of these non-cable bacteria species was below the detection limit of 1 pA/V, and more than 6 orders of magnitude lower than the average conductivity in cable bacteria (Table S1).

**Figure 1.**
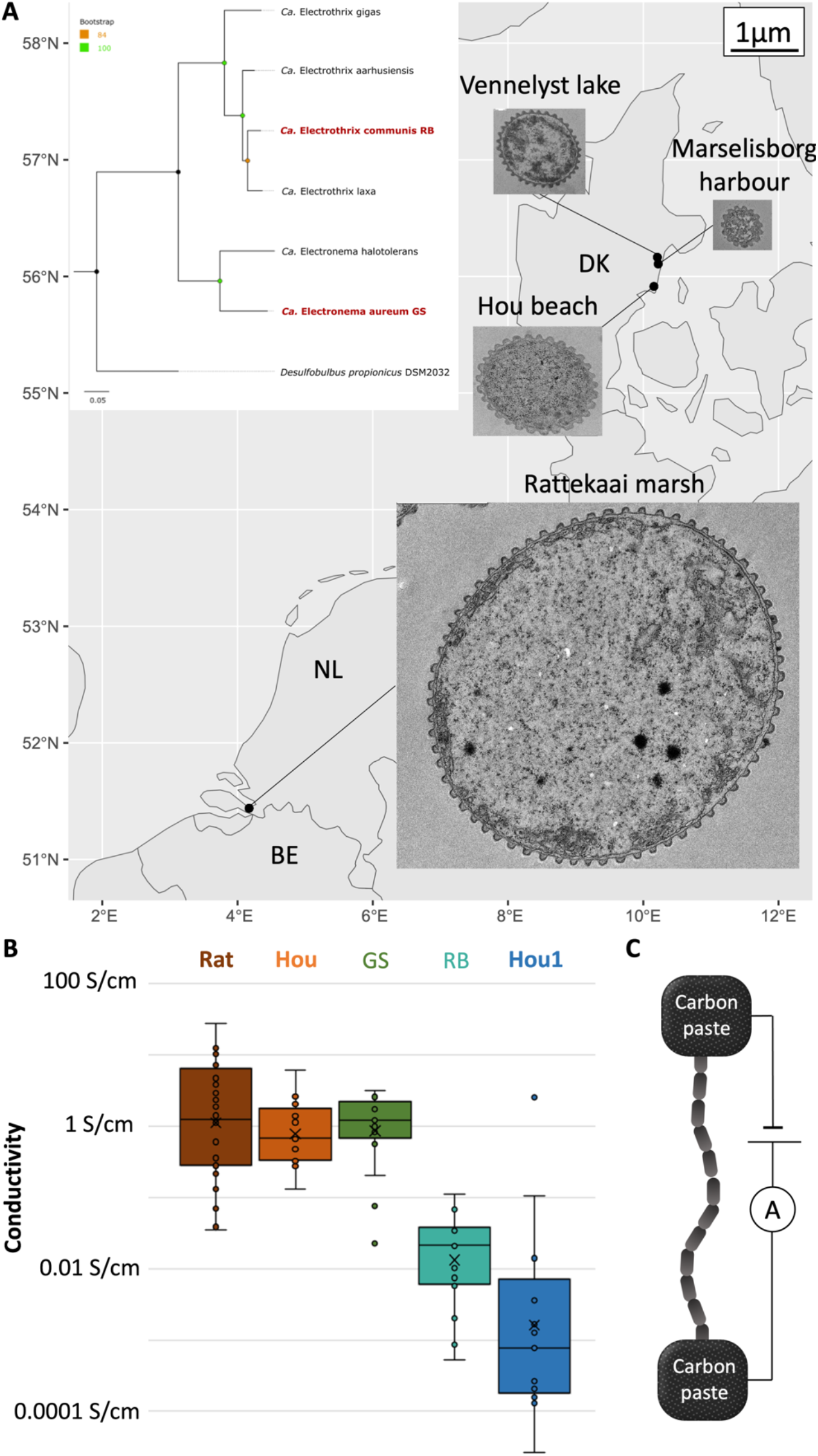
**A:** Geographic locations of samplings sites of cable bacteria investigated in this study and representative TEM cross sections from of cells from individual sites showing differences in cell diameter and number of ridges. Top right corner shows the scale bar for the TEM images. Panel insert: Phylogenetic tree of cable bacteria with the two enrichment cultures used in this study outlined in red. Scale bar shows the number of amino acid changes per site. **B:** Electrical conductivity of different cable bacteria strains (calculated for a single PCF 33 nm in diameter as measured in this study – see below). **C:** Cartoon of the experimental set-up for conductivity measurements of filamentous bacteria.

Overall current density measured for an entire cable bacterium filament is ultimately channeled through the PCFs, i.e. their combined cross-sectional area, and not the whole area of the filament. Furthermore, different strains of cable bacteria have different number of PCF as a function of their cell diameters (Cornelissen *et al*. 2018 and this work). Thus, we hypothesized that the conductivity of the individual PCFs in cable bacteria is highly similar, and thicker bacterial filaments are simply forming more PCFs to adjust to their higher metabolic needs. To assess and compare conductivity of different cable bacterial species, conductivity measurements were carried out under standardized conditions with a fixed gap length between the electrodes, measurement time and extraction time (Figure 1C). The only characteristic that might have affected the conductivity measurements except for the sample preparation was the physiological state of the extracted bacterial filaments. We suggest that the relatively high variability in electrical conductivity within the species presented below can be attributed to the yet unexplored factors, such as the cable bacteria activity at the time of sampling, cell division events and bacteriophage infections.

The electrical conductivity of the PCFs from different cable bacteria strains were shown to be relatively similar for the GS, Hou and Rattekaai samples (Figure 1B) with average values of 1,4 ± 0,9 S/cm, 1,3 ± 1,4 S/cm, and 4,2 ± 6,0 S/cm respectively. The highest conductivity of 27,5 S/cm was measured for a single Rattekaai cable bacterium. In comparison to the other strains RB and Hou1 cable bacteria showed surprisingly low conductivity of 0,03 ± 0,03 S/cm and 0,02 ± 0,07 S/cm, respectively. The values presented here were calculated for a single PCF with a diameter of 33 nm (as measured in this study, see below). All strains tested show linear current/voltage curves and equal stability in nitrogen atmosphere (Figure S1). In air, however, conductivity quickly decreases (Figure S1) as previously observed (Meysman *et al*. 2019).

### Nickel within the periplasmic conductive fibers

A unique elemental profile had previously been observed in a single cable bacteria strain which resulted in a model where nickel was suggested to be important for the conduction of cable bacteria from Rattekaai sediment (Boschker *et al*. 2021; Thiruvallur Eachambadi *et al*. 2021). To investigate if this is a universal property of cable bacteria and increase the resolution related to ultrastructural localization of the nickel ions, we employed two complementary elemental analysis tools on different cable bacteria strains. We compared the cellular presence and localization of metal ions for two single strain enrichment cultures (GS and RB) of cable bacteria. In both strains nickel was repeatedly detected by both methods in good agreement with the previously published data on an unidentified marine species (Boschker *et al*. 2021). Our samples of intact cable bacteria and extracted PCFs were analyzed in parallel using STEM-EDX. The nickel signals were present in spectra of both intact cells and extracted skeletons, indicating that nickel was localized in the PCFs in the periplasm and not in the cytoplasm or in membrane-bound proteins (Figure 2A). The raw signal count for iron, which is a common element in metalloproteins, was higher than the nickel signal in intact cable bacteria. However, after the skeleton extraction procedure the iron signal significantly decreased (Figure 2A).

**Figure 2.**
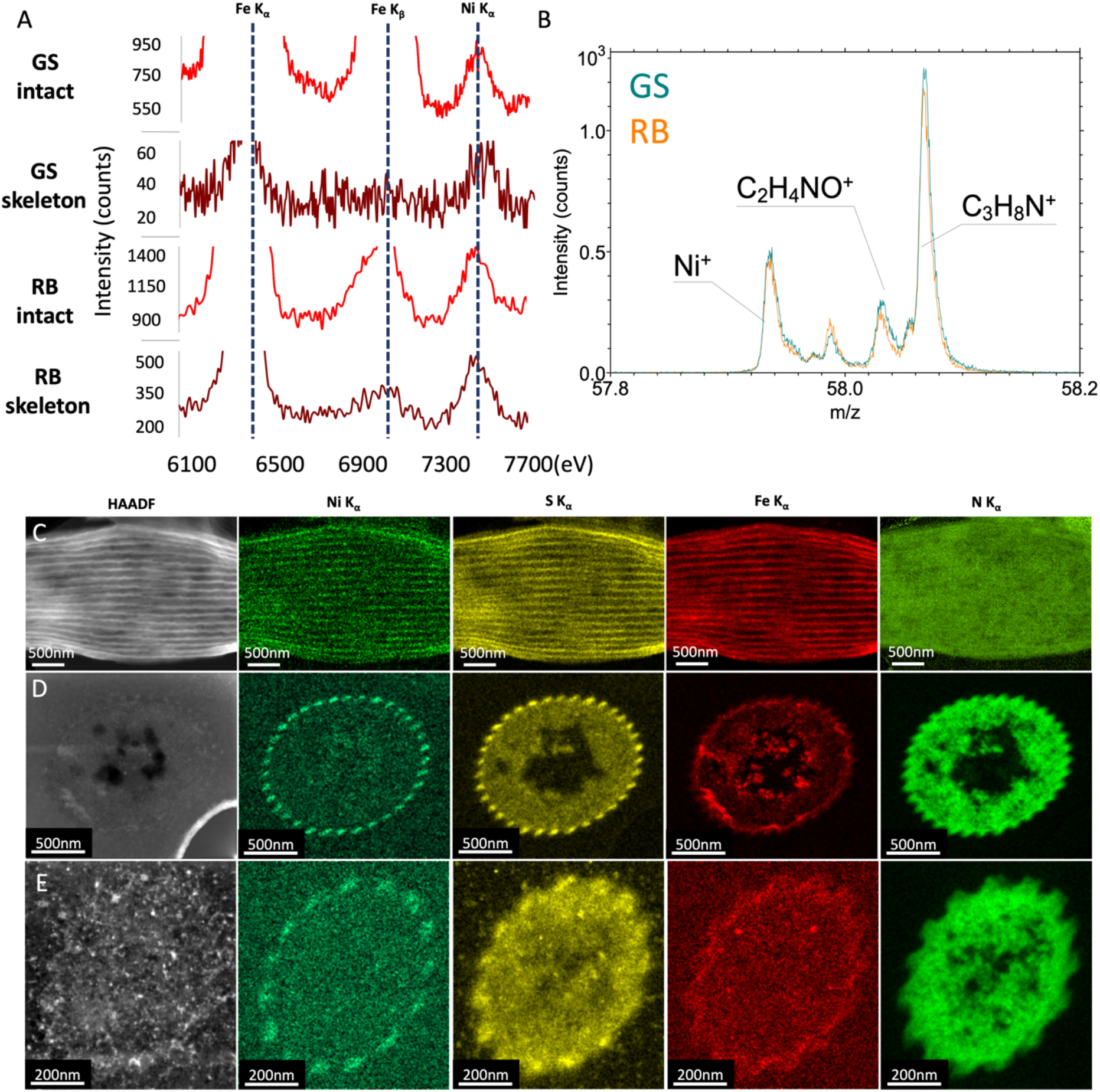
**A:** STEM-EDX spectra of single intact filaments and PCF-skeletons of GS and RB cable bacteria. Selected characteristic X-ray energies are shown for iron and nickel. **B:** ToF-SIMS spectrum showing presence of nickel in both GS and RB intact cable bacteria (represented by the cyan and orange curve, respectively). **C:** HAADF image of an untreated GS cable bacterium cell and STEM-EDX images showing the distribution of nickel, sulfur, iron and nitrogen. **D:** HAADF image of one 80 nm thick cross-section of a unstained GS cable bacterium cell and STEM-EDX images showing the distribution of nickel, sulfur, iron and nitrogen. **E:** HAADF image of one 80 nm thick cross-section of a unstained RB cable bacterium cell and STEM-EDX images showing the distribution of nickel, sulfur, iron and nitrogen.

In addition, isolated intact cellular filaments of both RB and GS were identically prepared and analyzed using the same ToF-SIMS equipment located at IMEC-Leuven (Belgium) as used in the earlier study on the Rattekaai marine cable bacteria (Boschker *et al*. 2021). The spectra of both strains clearly show the presence of signature nickel peaks (Figure 2B), which were further confirmed by isotope analysis (Figure S2). Moreover, the ToF-SIMS spectra of both RB and GS filaments exhibit numerous similarities (Figure S3) and display the characteristic protein and carbohydrate signatures that have previously been associated with the core-shell structure and basal sheath of the PCF (Boschker *et al*. 2021).

Since the conductive core was hypothesized to be made of protein with Ni-S co-factors, we then looked for potential co-localization of sulfur and nickel peaks in intact cells and extracted skeletons and their potential co-localization. STEM-EDX of intact cable bacteria detected a significant sulfur signal aligned along the axis of the PCFs, as also observed for nickel and iron (Figure 2C). To confirm that nickel and sulfur did indeed colocalize, principal component analysis (PCA) was performed in combination with non-negative matrix factorization (NMF) (Figure S4). Indeed, nickel and sulfur co-localize at the expected position of the PCF. Based on the Ni, S and Fe signals, measured PCF thickness was approximately 33 nm, in concordance with measurements from cross sections (Figure 2C). Finally, elemental analysis of cellular cross-sections of both species confirmed the presence of nickel and sulfur as well as a significant iron signal not described before in the PCFs (Figure 2D and E). Raw summed spectra of these samples can be found in Figure S5.

### Cross-sectional TEM analysis of intact cable bacteria filaments

No systematic comparative study of cable bacteria cell morphology and ultrastructure based on phylogenetically distinct species has previously been conducted. We collected high resolution TEM images of cross sections from cable bacteria strains of different phylogenetic origin embedded in plastic (GS and RB). The diameter of the PCFs of GS was measured to be 32,82 ± 3,22 nm (n = 34). The PCFs were equally spaced by an average distance of 107,75 ± 13,29 nm (n = 34). In the region between individual ridges the periplasm occupied a very limited space between the two inner and outer membranes and in most cases, it was not possible to reliably measure its thickness (Figure 3C and Figure S6C, D). The morphology of the PCF in RB was more rectangular, with a few exceptions (Figure S6A, B). Thus, we measured the height and width of the PCFs which were 25,95 ± 6,47 nm and 43,01 ± 13,5 nm respectively. Surprisingly, the cross-sectional area of a single PCF in RB, measuring approximately 1116 nm^2^, was around 50% bigger than that of GS, measuring around 846 nm^2^. Although, the total surface of all PCFs combined in RB is 40% smaller than that of the GS: 16 741 nm^2^ vs. 28 764 nm^2^(Figure S6)

**Figure 3.**
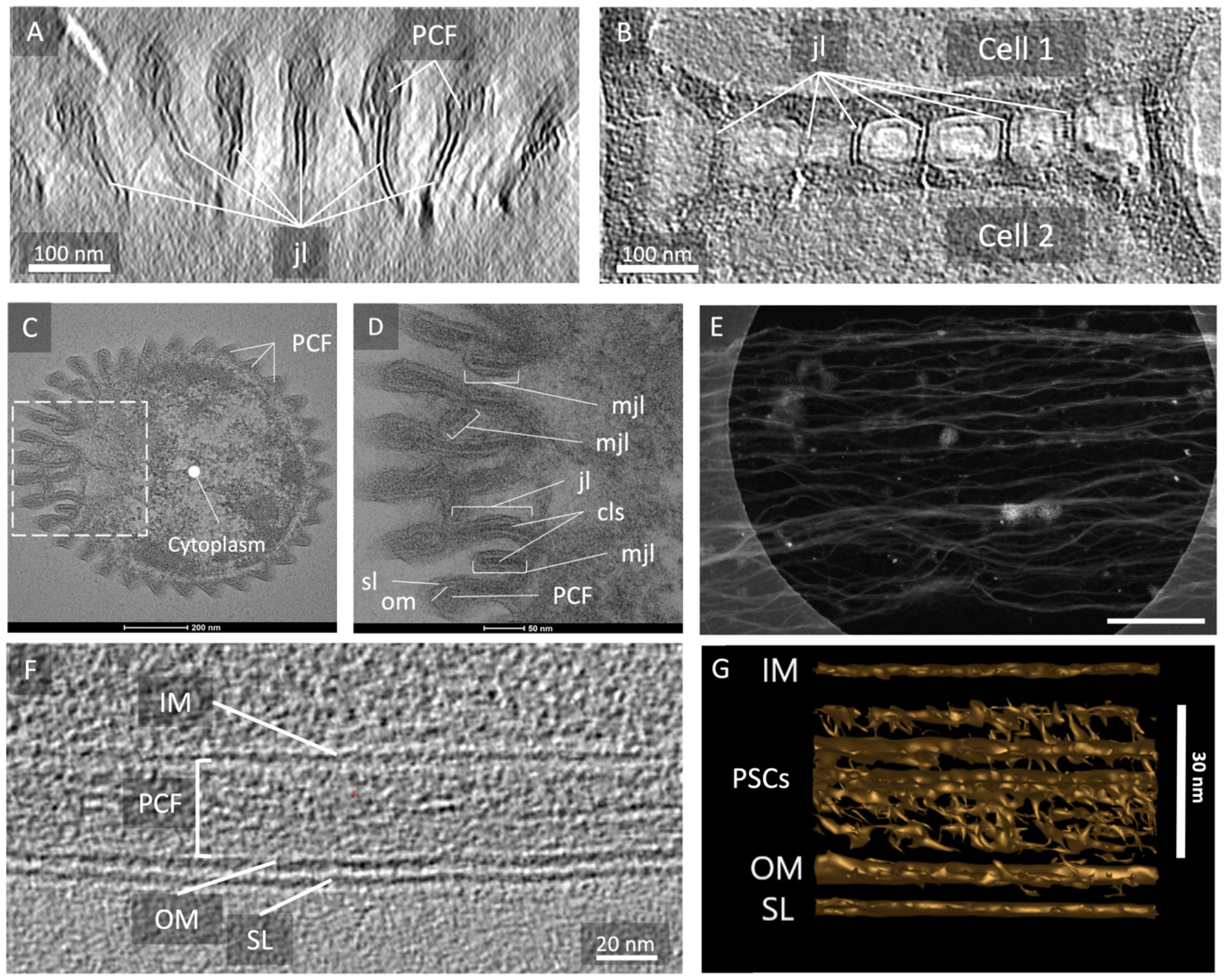
A: Cryo-ET cross-sectional view of cable bacteria cell-cell junction area. B: Cryo-ET longitudinal section view of GS cell-cell junction lamellae. C: Cross-sectional TEM image of intact cable bacterium close to the cell junction. Dashed square is the area zoomed in D. D: TEM image of a cross section showing cable bacteria cell junction and envelope compartments. E: HAADF image of PSC extracted from cable bacteria. Scale bar = 400 nm. F. Cryo-ET longitudinal section of intact cable bacterium showing the internal structure of one surface ridge. G. Sub-tomogram average of the cell structures in F. SL – surface layer, om – outer membrane, jl – junction lamella structure, mjl – minor junction lamella, PSC – PCF strand components, CLS – core lamella sheet.

In plastic-embedded cross sections from multiple cable bacteria species (Figure 3C, D and Figure S6) the surface of cable bacteria was observed to be composed of a surface layer (S-layer) that has not been described before. Particularly interesting was the morphology of the cell-cell junction region (Figure 3A, B, C, D). From each PCF towards the middle of the cell’s junction area a lamella-like structure with a thickness of 26,37 ± 1,44 nm (n = 10) was observed. This ultrastructural region was composed of the outer membrane having a thickness of 5,13 ± 0,75 nm (n = 10) and the S-layer with a thickness of 3,82 ± 0,66 nm (n = 10) on both sides of the lamella with an unknown internal structure showing high order in the center (Figure 3B) with a thickness of 8,0 ± 0,6 nm (n = 4). Overall, this ultrastructural lamella component is hereafter termed junction lamella (Figure 3A, B, D). Our data, thus, confirmed that the cell junction contains more than just two folded membranes (Cornelissen *et al*. 2018). Thiruvallur Eachambadi et al. previously hypothesised the existence of an interconnected structure at the cell junction important for the electrical robustness of the internal conductive network (Thiruvallur Eachambadi *et al*. 2020). Intriguingly, the lamellae seemed to be directly connected to the PCFs. Moreover, we observed an unusual structure in multiple cell junction cross sections, which appeared as a minor cell junction lamella lacking its PCF at the tip, but still containing the outer membrane fold and the core lamella sheet (CLS) (Figure 3D).

### Strand components of the periplasmic conductive fibers

Cable bacteria filaments completely lost their integrity after harsh mechanical manipulation (Figure S5) or being exposed to prolonged SDS treatment (Figure 3E). On the high-angle annular dark-field (HAADF) image shown in Figure 3E it was seen that after the extended SDS treatment the cytoplasm was removed as well as the carbohydrate layer that was presumed to keep the PCFs aligned in parallel (Boschker *et al*. 2021). Moreover, the 33 nm PCFs themselves appear unwound, falling apart into thinner strands. We termed these PCF strand components (PSC) and in some areas bundles of multiple PSCs were observed with the smallest single strands being 3,1 ± 0,2 nm (n = 17) in diameter. Such strands were consistently found in samples of different cable bacteria strains using SEM and TEM (Figure S7).

Cryo-ET of intact cable bacteria filaments also indicated that the 33 nm PCF contained PSC (Figure 3F, G). Sub-tomogram averaging of the longitudinal cross-section through the surface ridge structure revealed 3D volumes of the inner membrane, a PCF with multiple strand components, an outer membrane as well as the S-layer (Figure 3G). Surprisingly, the PCS did not seem to form a tight, crystalline superstructure but displayed a loose composition in the PCF.

### CryoET of intact cable bacteria filaments

After rapid fixation by plunge freezing in liquid ethane cable bacteria that were thicker than 600nm in diameter (GS and Hou) were milled with a Focused Ion Beam (FIB) in a Scanning Electron Microscope to a thickness of approximately 250 nm, which allows for collection of cryo-ET data of a cellular volume slice with good contrast. Multiple tilt-series for cryo-ET were collected from the resulting “windows” into the cells. This approach allowed us to see the morphological details in intact – non-stained – cable bacteria filaments in a frozen hydrated state and revealed a number of novel cellular structures described below.

#### Cytoplasmic filaments and tubes

Both GS cable bacteria and cable bacteria originating from environmental samples from Hou beach contained unknown filamentous structures with diameters between 7 nm and 28 nm in their cytoplasm. Some of these unidentified filaments (Figure 4A, B) displayed repetitive helical structure and formed straight parallel bundles. Other filaments (Figure 4C) were thin and curved. One kind of elongated structures were hollow (outer diameter 16 nm, inner diameter 11 nm) and we termed these nanotubes (Figure 4D, E). They appeared mostly empty, straight and with a less dense area in the center (Figure 4E). In some samples, dark filaments were observed (Figure 4D, E), with a more homogeneous, ordered structure. For some cells only one kind of filament was observed whereas for other cells multiple kinds of filaments were observed together in the cytoplasm. Interestingly, no such cytoplasmic filaments were observed in RB, even though the cryo-ET data in their case was collected on whole intact cells without FIB-milling.

**Figure 4.**
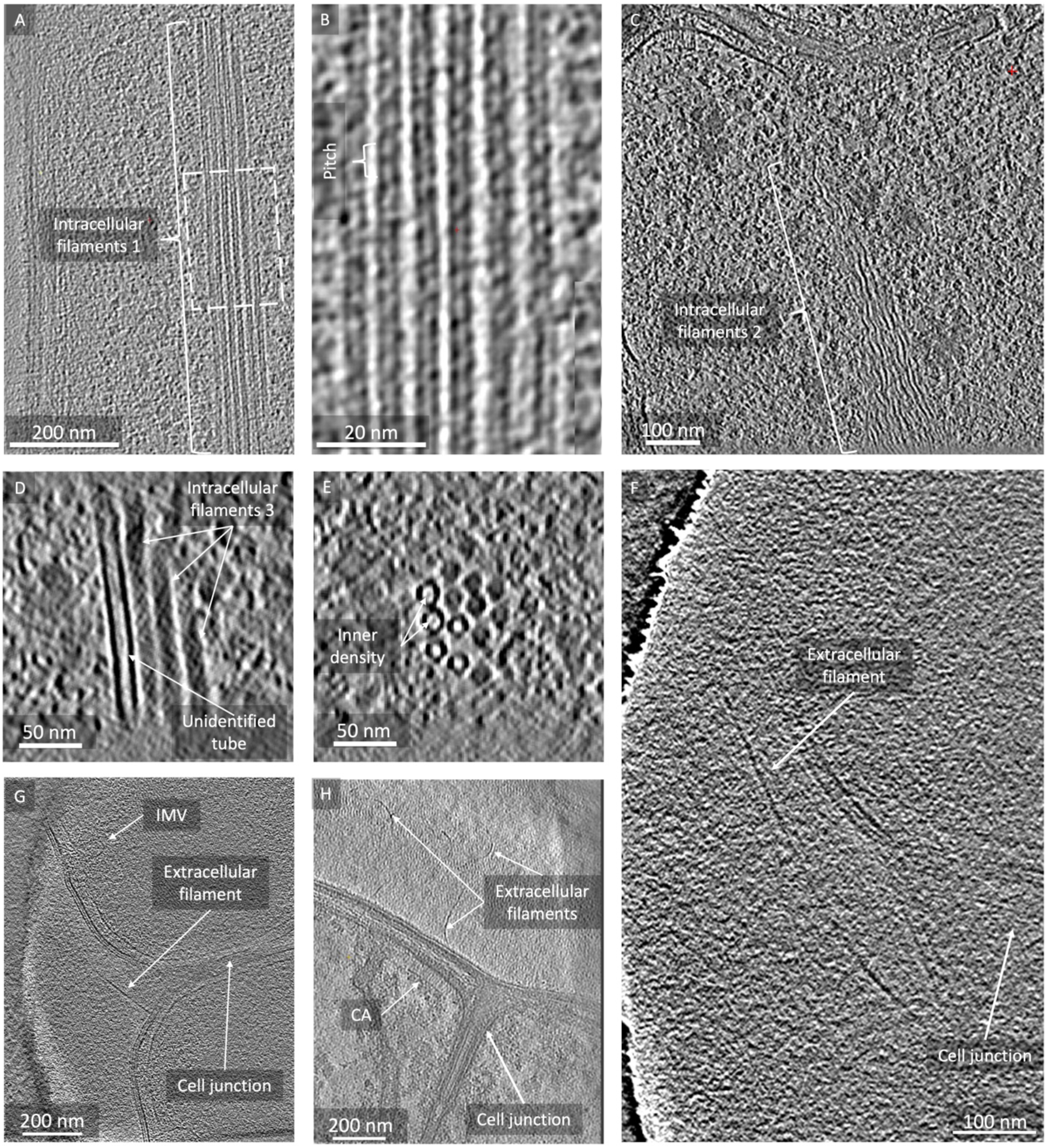
A: Cryo-ET of a GS cable bacterium with unidentified helical filaments in the middle of the cytoplasm Dashed rectangle is the area shown in B. B: Close-up of the helical filaments. C: Cryo-ET showing curved filaments inside of a GS cable bacterium. D, E: CryoET showing longitudinal and cross sections of the tubular and dark filaments inside of a GS cable bacterium. F: Longitudinal section of an RB cable bacterium with a long extracellular filament originating from the junction. G: Cryo-ET of the cell junction area from RB cable bacterium with a short extracellular filament originating from the junction. H: Cryo-ET of a Hou cable bacterium with multiple extracellular filaments close to the cell junction and a chemosensory array (CA).

#### Extracellular filaments

CryoET of RB and marine cable bacteria from Hou beach revealed extracellular filaments with a diameter of 5,92 ± 1,02 nm protruding from the cell-cell junctions. Morphologically they resemble type IV pili (Figure 4F, G) (Craig *et al*. 2003). The extracellular filaments were protruding from a region of cell in close proximity to the observed chemosensory arrays (Figure 4H).

#### Chemosensory arrays

Another type of cellular structures that was observed in both GS, RB, and in cable bacteria sampled from Hou beach were chemosensory arrays. Multiple inner membrane-bound chemosensory arrays were found at the cell poles (Figure 4H, and Figure 5A), docked either to the sides of or facing the cell-cell junction. In a few cases, however, the arrays were observed closer the middle of the cell, which could coincide with the start of a cell division event. In Figure 5C the general architecture of a cable bacteria chemosensory array can be seen. On average the arrays extended 35,8 ± 1,5 nm (n=16) into the cytoplasm of GS.

**Figure 5.**
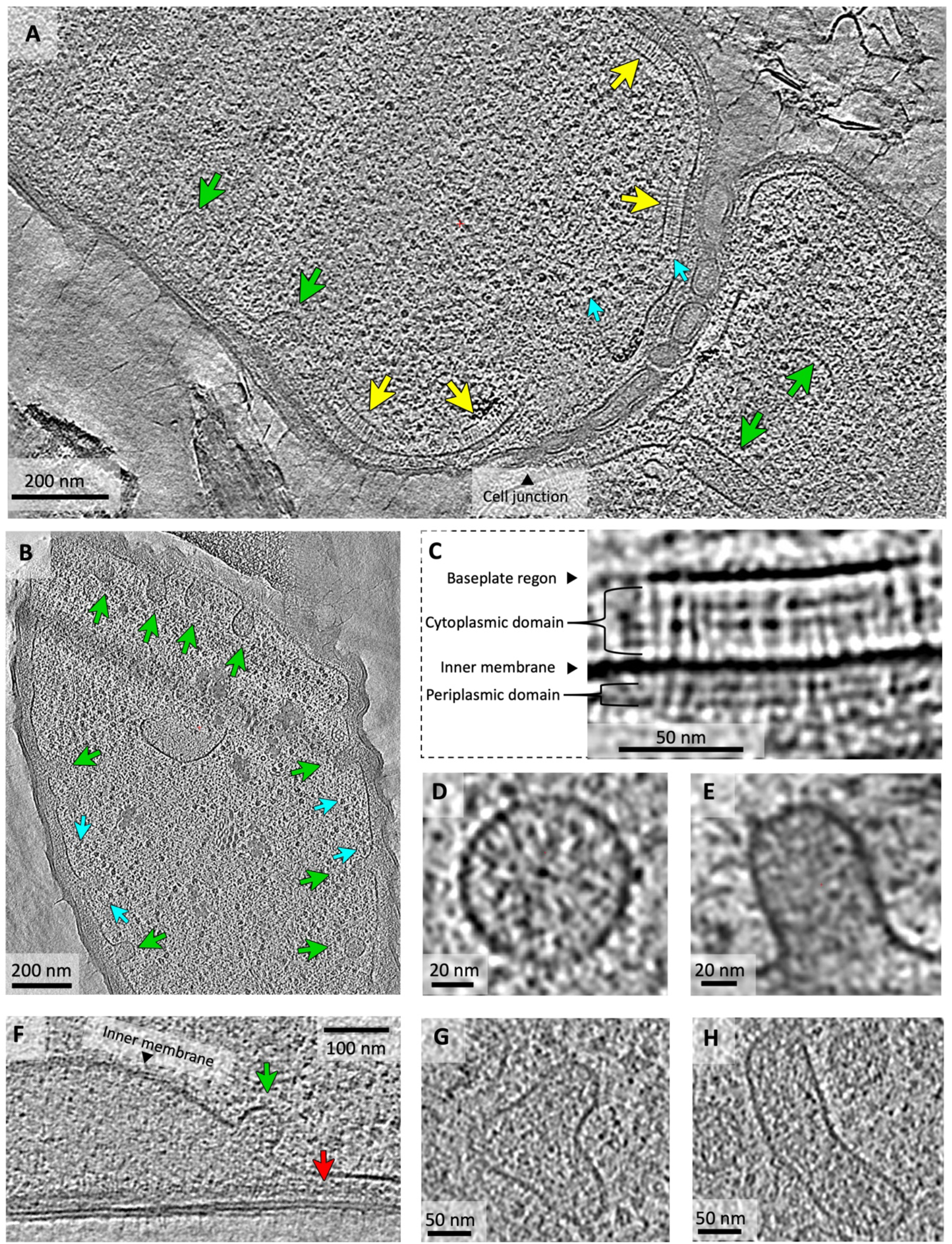
A: Cryo-ET of a longitudinal section of GS cable bacterium with multiple chemosensory arrays close to the cell junction. B: Cryo-ET of almost perfect cross section of GS cable bacterium displaying ridge structures and multiple cytoplasmic vesicles fusing with the periplasm. C: Close-up of a single chemosensory array with its main domains: base plate region, cytoplasmic domain and periplasmic domain. D: Inner Membrane-attached Vesicles (IMVs) top view. E: IMV side view. F: Inner membrane bulging event. G-H. Close-ups of cytoplasmic vesicles fusing with each other. Arrows: yellow – chemosensory array, cyan – putative polysomes, green – cytoplasmic vesicles, red – PCF.

#### Inner Membrane-attached Vesicles

In every cable bacteria cell investigated with CryoET we found multiple cytoplasmic vesicle-like structures (Figure 4G and Figure 5A, B). The vast majority of the vesicles were frozen at the moment of fusion with the inner membrane and we therefore termed them Inner Membrane-attached Vesicles (IMVs) (Figure 4G and Figure 5B, E). A few vesicles, however, were still found free in the cytoplasm, either singular or fusing with each other (Figure 5D, G, H). These free vesicles comprised less than 10% of the total vesicle count in our cryo-ET data, with the majority of the vesicles being IMVs. In a few tomograms inner membrane bulging was observed (Figure 5F), which formed a dome-shaped structure extending towards the cytoplasm from the periplasm. The contents of the domes were visibly different from the cytoplasmic content and no large structures were observed in this periplasmic matrix. Noteworthy, cytoplasmic vesicles were observed to fuse with the dome structures, too, and the PCFs did not seem to be affected by the bulging event, meaning that they still appeared aligned with the ridge, as shown in Figure 5F.

## Discussion

We set out to examine the hypothesis that cable bacteria, despite a quite high apparent diversity among species, must share a number of important features related to their conductive properties including organization and composition of their conductive fibres (Sereika *et al*. 2023). Our comparative studies of *Ca*. Electronema aureum GS (GS) and *Ca*. Electrothrix communis RB (RB) including investigations of elemental distribution in the periplasm, PCF conductivity and their ultrastructural organization greatly supported this hypothesis and revealed new important details of the conductive machinery and related structural features such as the CLS and IMVs that warrants further investigations. Based on these new data we updated the structural model of cable bacteria filaments (Figure 6).

**Figure 6.**
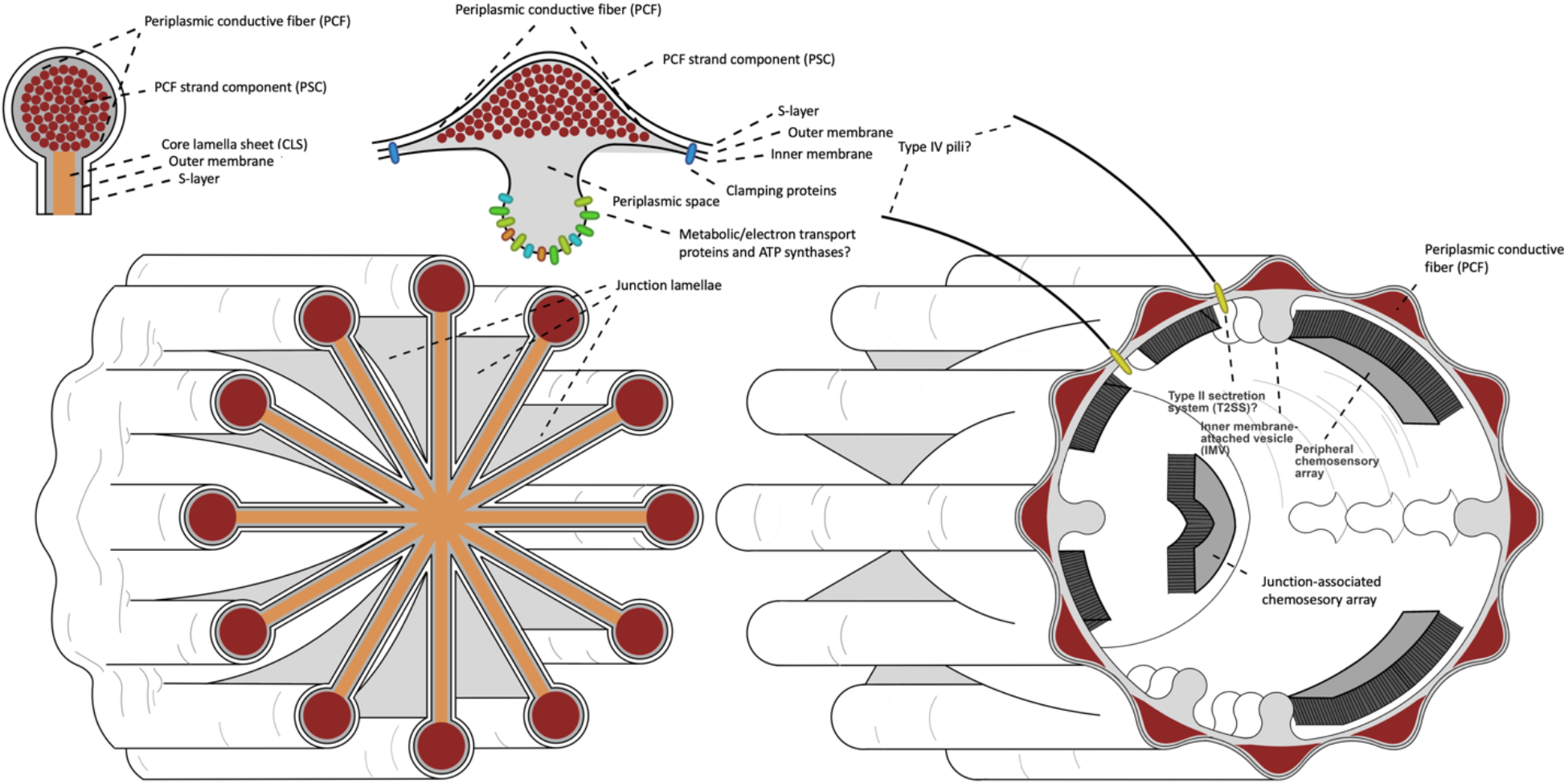
A new proposed morphological model of the cable bacteria cells. “?” designates hypothesized structures.

A highly localized nickel signal that clearly overlapped with a similarly significant sulfur signal in the periplasmic space of the characteristic ridge structures was shown with unprecedented detail. This clearly showed the ultrastructural organization of this element having a complete overlap with the PCF structure and underscored the potential importance of a nickel-sulfur co-factor in the conductive machinery of cable bacteria. The strength of the signal indicates that a substantial concentration of nickel could be present in the conductor, but unfortunately our data does not allow for a quantitative description of nickel in the ridge periplasm.

Our precise localization of nickel to the ridge periplasm as a part of the PCF in two distinct cable bacteria species and genera strongly supports that nickel is a universal component of the conductive fibers in cable bacteria, and it is conceivable that it plays an essential role in the long-range electron transport observed in cable bacteria. This finding carries significant implications for the underlying transport mechanism.

Previous studies have employed ToF-SIMS in conjunction with diverse analytical techniques to elucidate the composition of the conductive periplasmic fibers in marine cable bacteria (Boschker *et al*. 2021) retrieved from Rattekaai salt marsh. These combined efforts suggested a core-shell model, in which the fibers resemble a household electrical wire, featuring a conductive core surrounded by an insulating protein shell and resting on a basal sheath rich in polysaccharides. We did not see any evidence for an insulating protein-based structure around the PCF in our data.

In earlier studies, dissection of cable bacteria filaments by an AFM probe (Jiang *et al*. 2018) revealed dimeric structure of the periplasmic fibers in some of the preparations, with the diameter ranging between 10 and 30 nm. We show that the periplasmic fibers can be taken apart down to PCF strand components (PSC) level of approximately 3nm in diameter. In addition, our data shows a consistent diameter of the PCF in *Ca*. Eletronema aureum GS cable bacteria around 33 nm. Combined, this indicates that the periplasmic fiber is composed of multiple PSCs forming a loose stranded rope-like structure, and that the 10 nm fibers observed previously were probably not intact PCFs, but likely mechanically broken by the AFM tip into oligomers of PSCs.

Electrical conductivity measurements of the PSCs are challenged by difficulties in consistent production of these strands and is subject of ongoing studies. A great surprise was observed in the cryo-ET data, where the periplasmic fibers appear amorphous without any conspicuous crystallinity and this raises the question how the high conductivity observed for some cable bacteria can be obtained.

Considering the data presented here, we suggest that viewing PCFs as perfect cylinders is overly simplistic and the volume occupied by the PCF in the ridges is dynamic. Such architecture could allow for a fine tuning of electrical conductivity by expulsion of PSC building blocks from the putative biosynthesis machinery into the periplasm, where they would polymerize. Such a mechanism could explain construction of new PCFs for higher metabolic needs of thicker cable bacteria species.

Cable bacteria filaments have a different number of ridges containing periplasmic fibers, depending on the size (Cornelissen *et al*. 2018), but it is yet unknown by what mechanism they are formed. We speculate that the biosynthesis machinery involved in producing PSCs and expanding PCFs could be located at the septum when cell division is initiated.

An intriguing observation form our high resolution ultrastructural studies was the revelation of the junction lamellae composition with a highly ordered core material – the CLS – that is likely the material that interconnects the fibers in the cell-cell junctions making the long-distance conduction fail-safe (Thiruvallur Eachambadi *et al*. 2020) (Figure 6). A biosynthesis machinery associated with septum formation would also allow for the association of PCFs and CLS in the cell-cell junction. The evidence of electrical conductivity in the CLS structure and the fact that it has a structural composition distinct from the PCF makes it a very interesting target for future investigations of structure, molecular composition, and conductive properties.

In 2019 PilA protein was shown to be the most abundant protein in cable bacteria Ca. Electronema aureum GS and since then its role has been extensively discussed (Kjeldsen *et al*. 2019). PilA was hypothesized to build the PCF enabling electron transport akin to e-pili of *Geobacter* bacteria.

In other microorganisms, PilA builds type IV pili that perform multiple functions like DNA capture and motility(Craig *et al*. 2019; Denise *et al*. 2019). For twitching and gliding motility type IV pili function as improvised grappling hooks. They extend in the direction of movement, attach to a surface and then retract moving the cell forward (Merz *et al*. 2000; Skerker and Berg 2001). In *Myxococcus xanthus* type IV pili are essential for what is called social gliding motility or S-motility(Kaiser 1979; Wu *et al*. 1997) which is controlled by a chemosensory system(Sun *et al*. 2000). During S-motility cells connect to each other and move as a pack. Such social gliding is usually accompanied with exopolysaccharide (EPS) or slime secretion(Li *et al*. 2003). Among filamentous bacteria *Beggiatoa, Thiothrix, Thioploca* (Larkin and Strohl 1983) and cyanobacteria(Adams 2001; Risser and Meeks 2013) glide on a trail of EPS, but the exact mechanisms there are not clear.

Cable bacteria, too, are capable of gliding motility (Bjerg *et al*. 2016), and they secrete EPS as they glide (Geerlings *et al*. 2019; Kjeldsen *et al*. 2019). The presence of putative type IV pili shown in this study suggests a motility mechanism by which cable bacteria propel themselves attaching to a surface with pili and gliding over EPS as the pili are being retracted. The newly found chemosensory systems then will mediate signal transduction and regulate the motility (Figure 6).

We identified multiple types of cytoplasmic filaments and the molecular basis and function if these filaments is still unclear. We note that similar observations of cytoplasmic filaments have been made for other bacteria where the functions could not be assigned (Oikonomou and Jensen 2021). We speculate that some of the observed filaments could be phage-based but this will have to be verified in future studies.

Formation of cytoplasmic vesicles originating from the inner membrane in gram-negative bacteria is a poorly understood phenomenon and observations are rare (Toyofuku *et al*. 2019). In *Bacillus subtilis* cytoplasmic vesicles are known to be associated with phage-induced cell death (Toyofuku *et al*. 2017). We, however, observed IMVs in cable bacteria cells that had no signs of phage infection. IMVs in cable bacteria are abundant and we hypothesize that they are playing an important role in general cable bacteria metabolism. Küsel and co-authors speculated that inner membrane vesicles found in *Acidiphilium cryptum* were produced to increase cellular contact area with electron acceptors (Küsel *et al*. 1999). A recent study in *Geobacter sulfurreducens* showed that in conditions with limited energy availability bacterial cells produced stacks of intracytoplasmic membranes to enhance membrane-bound processes (Howley *et al*. 2023). Taking into account that the vast majority of the vesicles that we observed in different cable bacteria were attached to the inner membrane we suggest that they are used as activity hotspots for membrane-dependent cellular processes associated with PCFs. We also suggest that the membrane of the IMVs may contains electron transport chain components. Multiple studies have shown that components of the electron transport chain can form supercomplexes and that these complexes can induce curvature in the membrane as is also the case for ATP synthase (Blum *et al*. 2019; Mühleip *et al*. 2023) (Figure 6). This could explain the fusion-like morphology of the IMVs.

## Materials and Methods

### Cable bacteria strains

Freshwater cable bacteria (GS) were taken from the single strain enrichment culture of *Candidatus* Electronema aureum GS, Aarhus, Denmark (56.164796, 10.207805) (Thorup *et al*. 2021). Marine cable bacteria (RB) were taken from the single strain enrichment culture of *Candidatus* Electrothrix communis RB originating from Marselisborg sediment, Aarhus, Denmark (56.136972, 10.210128). Wild type marine Hou cable bacteria originate from a mixed culture sampled at Hou beach, Hou, Denmark (55.915086, 10.256861). Wild type marine Rattekaai cable bacteria originate from a mixed culture sampled at Rattekaai salt marsh, Rilland, Netherlands (51.439167, 4.169722).

### Bacterial filament extraction

Clean cable bacteria filaments were obtained by placing a piece of the sediment under a stereomicroscope and fishing cable bacteria out of the sediment by a home-made glass hook. The filaments were washed in drops of clean filtered Milli-Q water to remove sand and dust particles.

To extract the periplasmic conductive fibers (PCF), the cable bacteria were washed as described above and then were moved to another droplet with 1% weight/volume SDS for 10 minutes, washed with Milli-Q water up to three times, treated with 1mM EDTA, pH8, for 10 minutes, washed again and air dried. These extracted and purified PCFs were lastly moved to any substrate for further investigations, for example, Electron Microscopy grids, silicon substrate for EDX or electrodes.

To produce PCF strand components (PSCs) from the PCF of cable bacteria alternative treatments were used. Clean cable bacteria filaments were placed on EM grids and 4uL of SDS were added on the grid to completely cover its surface. After 100 minutes, of incubation, the samples were washed with 100uL to 150uL of MilliQ. The grid was directly taken for analysis. Mechanical extraction of PSC structures included cycles of freezing in liquid nitrogen and centrifugation at 12000g.

### Conductivity measurements’

Conductivity of cable bacteria was measured using a 4200 Keithley-SCS (Keithley, Solon (Ohio), USA). Cable bacteria filaments were fished out of the sediment and washed in MilliQ water at least 6 times before being placed on a glass substrate. The filament was dried with nitrogen gas for 30 seconds, before carbon paste (EM-Tec C33, Micro to Nano, Haarlem, the Netherlands) was applied on both ends of the filament. All of this was conducted within a 5-minute timeframe. The gap size was between 100 and 800 μm and measured afterwards the with a light microscope. The prepared sample was placed in a probe stage, which was flushed with N_2_ twice before being placed in (low) vacuum. Then, Tungsten/Iridium probes were mounted on the carbon paste electrodes and voltage scans of -1 to 1 V were conducted, while the current was monitored. From this the conductivity was calculated using *σ =* (*I · l*)/ (*V · A*) with *I* the current at *V =* 100 *mV, l* the conductive channel, and *A* the cross-sectional area of the filament This last one was calculated from the diameter of the cable bacterium, assuming 15 fibers per μm of diameter, assuming one fiber to be 850 nm^2^ for Rat, Hou and GS and 1100 nm^2^ for RB.

All control samples were prepared by placing one to hundreds of filaments on an interdigitated Au electrode with 50μm interdistance (PW4XIDEAU50, Metrohm, Oviedo, Spain). The sample was placed in a vacuum atmosphere to prevent possible degradation and ensure adhesion of the filament to the electrode surface. The same system as described above was used to find the conductivity. Now I is the cumulative current from all 10 parallel current paths, l the conductive channel (50 μm), and A the cross-sectional area of the filaments (0.5 μm^2^). This last one is a conservative lower limit assuming 5 filaments with a similar cross-sectional area to those of cable bacteria. With currents never exceeding the pA range, conductivities were always found to be below 10^−7^ S/cm. In most cases, more filaments were applied, ending up in conductivities below 10^−8^ S/cm. This was repeated twice for every species (n = 3), with one negative control being a dried drop of the culture medium, which gave currents in the same pA range.

### STEM-EDX

STEM-EDX spectra of intact cable bacteria, PCF-skeletons, and cross-sections were recorded on FEI Talos FX200i field emission gun transmission electron microscope operated in scanning transmission mode. Individual spectra and element distribution maps were acquired within 20 and 560 minutes at 33,000-94,000 times magnification with a beam current of 0,472 to 7,48 nA and a beam energy of 200 keV and a dwell time of 20.0 μs. Velox (ThermoFisher Scientific) was used to collect the raw data and produce binned elemental maps and Hyperspy (GitHub project DOI: 10.5281/zenodo.7263263) was used to extract raw summed spectra and for PCA and NMF.

### ToF-SIMS

Multiple cleaned intact filaments of cable bacteria were deposited on a gold-coated silicon wafer. The filaments were then located by both fluorescence microscopy and AFM for subsequent ToF-SIMS analysis. ToF-SIMS data was acquired using the TOF.SIMS 5 device (IONTOF GmBH, Germany) located at Imec, Leuven (Belgium). We used the high current bunched mode for maximal mass resolution with a Bi3^+^ analysis beam (30 keV, current ∼0.35 pA, 100 × 100 μm^2^ area, 512 × 512 pixels). Existing factory settings were optimized to obtain maximal current. This setup is traditionally employed for surface spectrometry; however, despite being slower than a dedicated sputter gun, the Bi_3_^+^ analysis beam could still penetrate through the filaments, and this arrangement ensured the collection of all filament material.

Data analysis was performed using the SurfaceLab 7 software (IONTOF GmBH, Germany). All measurements were calibrated using C^+^, C_2_H_3_^+^, C_3_H_4_^+^, C_3_H_5_^+^, C_4_H_5_^+^ and Au^+^. Mass spectra were obtained by summing over all pixels in the lateral region of interest (ROI) and over all data points in the depth ROI: substrate regions were excluded from the lateral ROI to minimize substrate counts, and the first ten data points were removed from the depth ROI since these data points are related to surface transient. The relative count of isotope ^x^Ni was calculated as the ratio of its absolute count to the sum of the absolute counts of the three most abundant nickel isotopes ^58^Ni, ^60^Ni and ^62^Ni. The expected counts were calculated based on the natural abundances of ^58^Ni (68.08%), ^60^Ni (26.22%) and ^62^Ni (3.64%) (Figure 2S).

### Cross-sectional TEM

Cable bacteria filaments were fished from the sediment and washed in a water droplet to remove dust and sand particles. Clean bacteria were quickly injected into a small drop of agarose and dumped into the fixative 0,1 M sodium cacodylate buffer solution containing 2,5% formaldehyde and 2,5% glutaraldehyde, and stored in the refrigerator, for three days. Afterwards, the sample was rinsed twice with 0,1 M Sodium Cacodylate buffer for 10 min and 3 times rinsed with dH_2_O for 5 min. and then rinsed 2 times for 10 min. with the increasing concentrations of ethanol as follows 50%, 70%, 90%, and 99%. With 99% ethanol washing being 2 times for 15 min. The last washing solution of propylene dioxide was used 2 times for 15 min. The sample was then moved to the mixture of propylene dioxide 1:1 in EPON epoxy resin and left overnight under the fume hood. The agarose drops with the cable bacteria filaments, from the propylene dioxide/Epon mixture were put into pure Epon for 8 hours, then placed in a mold and left to cure at 60 ºC for at least 48 hours. The resulting Epon block was then put into the microtome for sectioning by cutting with a glass knife for an overview, while thin sections were cut with a diamond knife at an angle of 45º. Selected 50-60 nm sections (70-80 nm for STEM-EDX) were put on homemade copper EM grids with 400 mesh size and covered with 2% collodion solution and a film of carbon using carbon evaporator. Post stain á La Reynolds was used to prepare the sections for TEM data collection.

### FIB-SEM

Cryo-Focused-Ion-Beam Scanning Electron Microscopy was done on Aquilos 2 (ThermoFisher Scientific). Samples for milling were frozen on 2/2 200 gold mesh grids by plunge freezing in Leica EM GP2 or GP1 plunge freezer (Leica Microsystems, Germany) at 10°C and 90% relative humidity. No fiducial particles were added. Two layers of platinum were deposited on the grid to protect the sample and ensure conductivity. Lamellae were milled with gallium ion beam to the target thickness of 250nm.

### CryoET

Cable bacteria samples were prepared as described above and milled with FIB-SEM or in their inctact form in the case of RB. Tilt series were collected on Titan Krios (Thermo Fischer Scientific, USA), 300kV, equipped with Gatan K3 camera. The dose was set to 2 e/Å^2^ for data collection on lamellae or 2.3 e/Å^2^ for data collection on intact cable bacteria at magnification of 26,000 x. The estimated pixel size was 3.385 Å with the defocus values ranging from 3.00 to 10.00 μm. Tiltseries of cable bacteria lamellae were collected dose-symmetrically from -60 to 60 degrees with increments of 2 degrees. Movies were motion-corrected using MotionCor2 (Zheng *et al*. 2017). Tomograms were then reconstructed with eTomo as a part of IMOD 4.12.21 (Kremer *et al*. 1996) and CTF-deconvoluted using IsoNet (Liu *et al*. 2022). PEET 1.16.0 was used for sub-tomogram averaging on raw tomograms(Nicastro *et al*. 2006; Heumann *et al*. 2011).

### DNA extraction and sequencing

Sediment cores with the clonal strain of marine cable bacteria (RB) were grown for 2 weeks at 20°C in a closed box with water in the bottom before sampling. The upper layer (≈1 cm) was scooped up, put in a petri dish, and tufts of cable bacteria were dragged out and washed in autoclaved seawater. The tufts were collected in aliquots of 0.25 mL into eight G2 DNA/RNA enhancer bead tubes (Ampliqon, Odense, Denmark). DNA was extracted with the DNeasy PowerLyzer PowerSoil kit (Qiagen, Hilden, Germany), where the provided bead-tubes in the kit were exchanged for the G2 DNA/RNA enhancer bead tubes. The 8 samples were merged in the provided clean-up micro-column. The DNA sample was eluted twice in 40 μL, where the first elution was prepared further for long read sequencing, and the second elution was prepared using the Nextera DNA Library Preparation Kit (Illumina) and sequenced using the Illumina MiSeq kit v3 with 2×300 bp reads. The samples for nanopore sequencing were prepared as follows: The DNA sample was size selected with BluePippin pulsed field electrophoresis (Sage Science, Beverly, MA, USA) in the range 2,000-50,000 bp and eluted in 60 μL. The sample was prepped for nanopore sequencing using the Genomic DNA by ligation (SQK-LSK110) protocol and sequenced by MINion (Oxford Nanopore Technologies, Oxford, UK) with the software MinKNOW (v.22.12.7, Oxford Nanopore Technologies, Oxford, UK).

### Genome assembly

The long reads were trimmed with Porechop (v.0.2.3_seqan2.1.1, (Wick *et al*. 2017)) and the metagenome assembled with Flye (v.2.8.1-b1676, (Kolmogorov *et al*. 2019)). The short reads were trimmed and paired with Trimmomatic (v.0.39, (Bolger *et al*. 2014)), and these were mapped to the metagenome and used to polish the metagenome 4 times with the software Racon (v.1.4.20, (Vaser *et al*. 2017)). All reads were mapped to the polished metagenome by Minimap2 (v.2.17-r941, (Li 2018)) and Samtools (v.1.9, (Danecek *et al*. 2021)), a depth matrix for the mapped reads were produced by jgi_summarize_bam_contig_depths (v.2.15) with a minimum end-to-end identity of 95% for each read, and the metagenome was binned into individual genome bins by Metabat2 (v. 2:2.15, (Kang *et al*. 2019)). The bin identified as a cable bacteria genome was analyzed with fastANI (v.1.33, (Jain *et al*. 2018)), Quast (v. 5.2.0, (Gurevich *et al*. 2013)) and CheckM2 (v.1.0.1, (Chklovski *et al*. 2022)).

### Phylogenetic Analysis

Published genomes of cable bacteria were collected along with the genome of *Ca*. Electrothrix communis RB. The 120 conserved bacterial single copy genes (Parks *et al*. 2018) were identified with the GTDB-Tk tool (v. 2.2.3, (Chaumeil *et al*. 2020)) and the contig count was analyzed with Quast. The genomes for the phylogenetic analysis were chosen on the basis of a contig count <50 and a minimum of 110 single copy genes present in the genome. These cut-offs were chosen to maximize the accuracy of the tree. The single copy genes were aligned with the GTDB-Tk and a phylogenetic tree was made with IQ-tree (v. 1.6.12, (Nguyen *et al*. 2015)) with 1000 Bootstraps and *Desulfobulbus propionicus DSM* as outgroup.

## Ancknowledgments

We want to thank for excellent technical support Pia B. Jensen, Taner Drace, Andreas Bøggild and Jesper L. Wulff. We thank Federico Aulenta of Italian National Research Council who provided us with three pure cultures of filamentous bacteria for conductivity measurements. We also thank Nikoline Sanggård Madsen, who brought our attention to polysomes in CryoET data, Xuya Yu, Mingdong Dong who assisted with probe stage conductivity measurements, Lars Damgaard and Jesper J. Bjerg who participated in project discussions, and, finally, Robin Bonné, who organized series of meetings between Aarhus and Hasselt Universities.

## Author contributions

Conceptualization: LD, MLJ, RB, NF, KW, EDB, LPN, JM, TB

Methodology: LD, MLJ, RB, NF, IM, JLH, EDB, JM, TB

Investigation: LD, MLJ, RB, NF, KW, PBJ, LEPJ, LNA

Visualization: LD, MLJ, RB, NF, KW, PBJ, LEPJ, TD, AB

Supervision: IM, JLH, AS, EDB, LPN, JM, TB

Writing – original draft: LD

Writing – review & editing: all authors

