## Supplementary data for "Comparative electric and ultrastructural studies of cable bacteria reveal new components of conduction machinery"

**This PDF includes:**
Supplementary Table S1
Supplementary Figures S1 to S7

**Table S1. Electrical conductivity measurements of different non-conductive filamentous bacteria.**

| | Reference/Source | Species/Genus | Conductivity (S/cm) |
|---|---|---|---|
| **Pure cultures** | Aulenta et al., Rome University | *DYN-VER9-ISO2* related to *K. aurantiaca* | < $10^{-7}$ |
| | Aulenta et al., Rome University | *Ca.* Magenema Perideroedes | < $10^{-7}$ |
| | Aulenta et al., Rome University | New genus within *Cytophaga flexibacter* | < $10^{-7}$ |
| | The Leibniz Institute DSMZ (5205) | *Thiothrix nivea* | < $10^{-7}$ |
| | The Leibniz Institute DSMZ (14523) | *Anaerolinea thermophila* | < $10^{-7}$ |
| | The Leibniz Institute DSMZ (16556) | *Leptolinea tardivitalis* | < $10^{-7}$ |
| | The Leibniz Institute DSMZ (23923) | *Pelolinea submarina* | < $10^{-7}$ |
| | The Leibniz Institute DSMZ (14018) | *Geothrix fermentans* | < $10^{-7}$ |
| | The Leibniz Institute DSMZ (21853) | *Caldiserica exile* | < $10^{-7}$ |
| | Kawaichi et al., (Kawaichi *et al.* 2018) | *Ardenticatena maritima* | < $10^{-7}$ |
| | Chailakhyan et al., (Chailakhyan *et al.* 1982) | *Phormidium unicatum* | < $10^{-7}$ |
| | Pasteur Culture Collection for Cyanobacteria (PCC) | *Geitlerinema PCC 9228* | < $10^{-7}$ |
| **Field samples** | Ferry Harbour, Grenå | *Beggiatoa* | < $10^{-7}$ |
| | Schleswig sediment | *Crenothrix* | < $10^{-7}$ |
| | Aggersund | *Synecoccus* | < $10^{-7}$ |

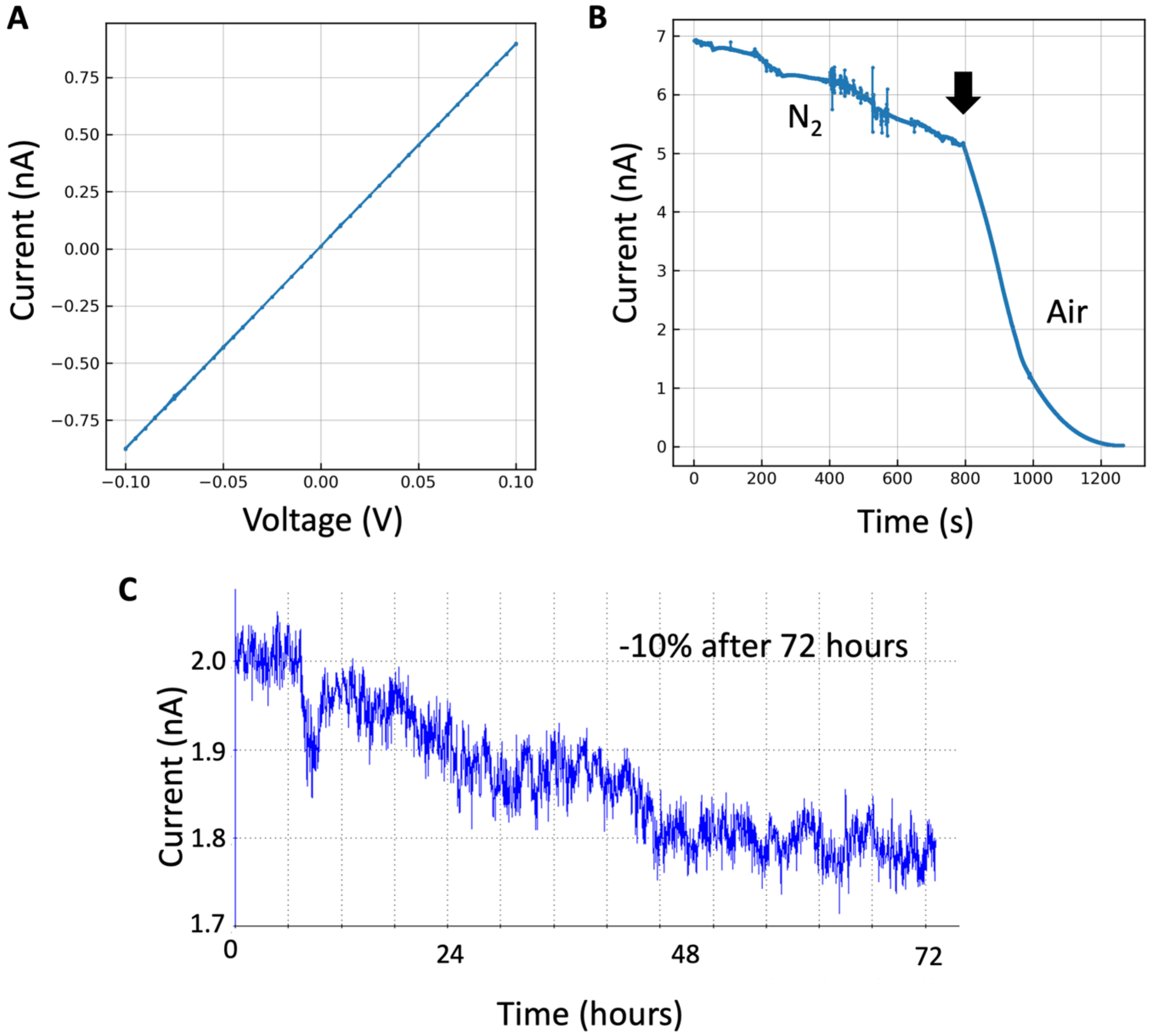


**Figure S1. Conductivity characteristics were shared among all tested cable bacteria strain**. **A:** the highly linear current response to the voltage; **B:** the slow degradation in a $N_2$ atmosphere, and fast degradation in air (black arrow shows when air was introduced); **C:** the stability of electrical conductivity in a vacuum at 100mV.

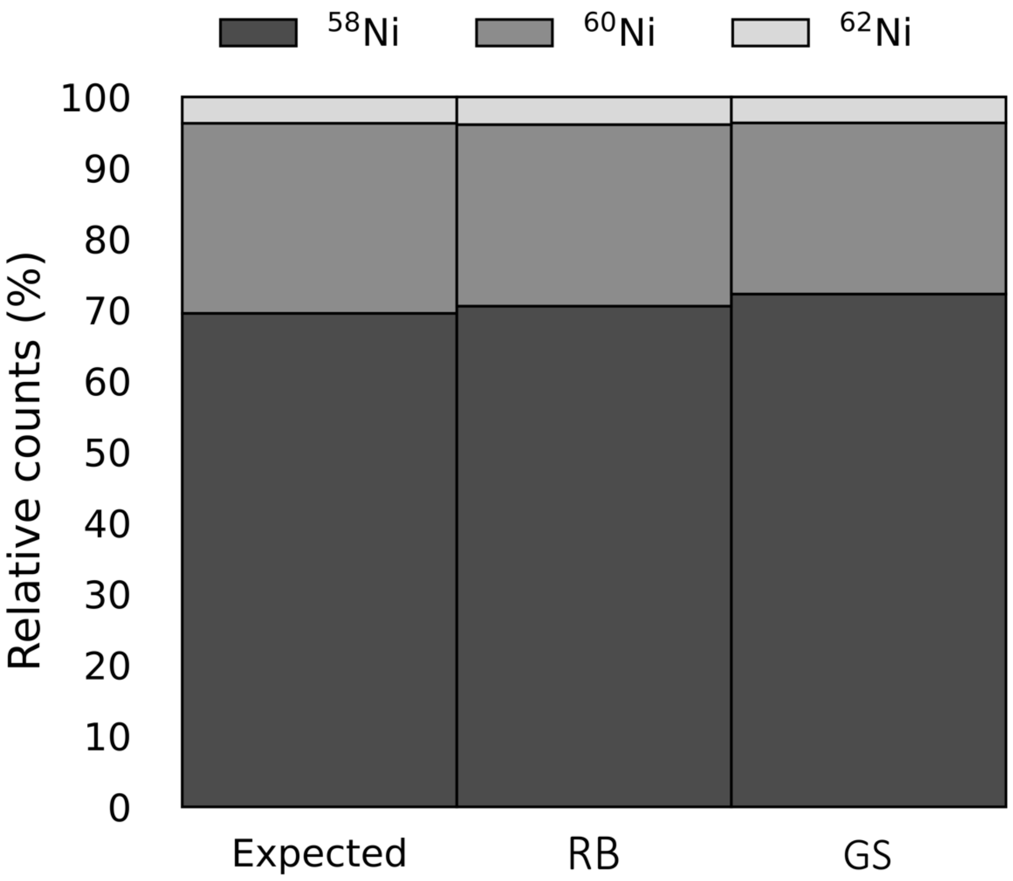


**Figure S2**: Isotope analysis of nickel for both strains, RB and GS, demonstrates agreement between the observed and expected counts, indicating that nickel is correctly identified.

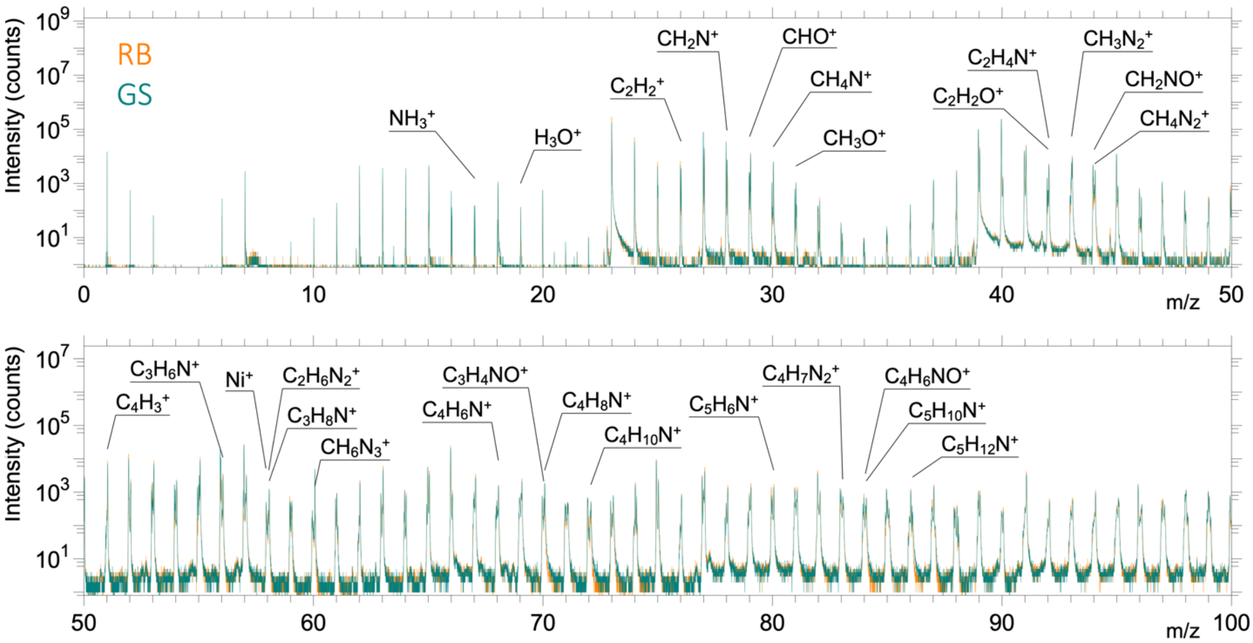


**Figure S3:** The ToF-SIMS spectra of intact RB and GS filaments exhibit similarities. The annotated fragments have been associated to protein and polysaccharide layers in previous studies (Boschker *et al.* 2021; Thiruvallur Eachambadi *et al.* 2021).

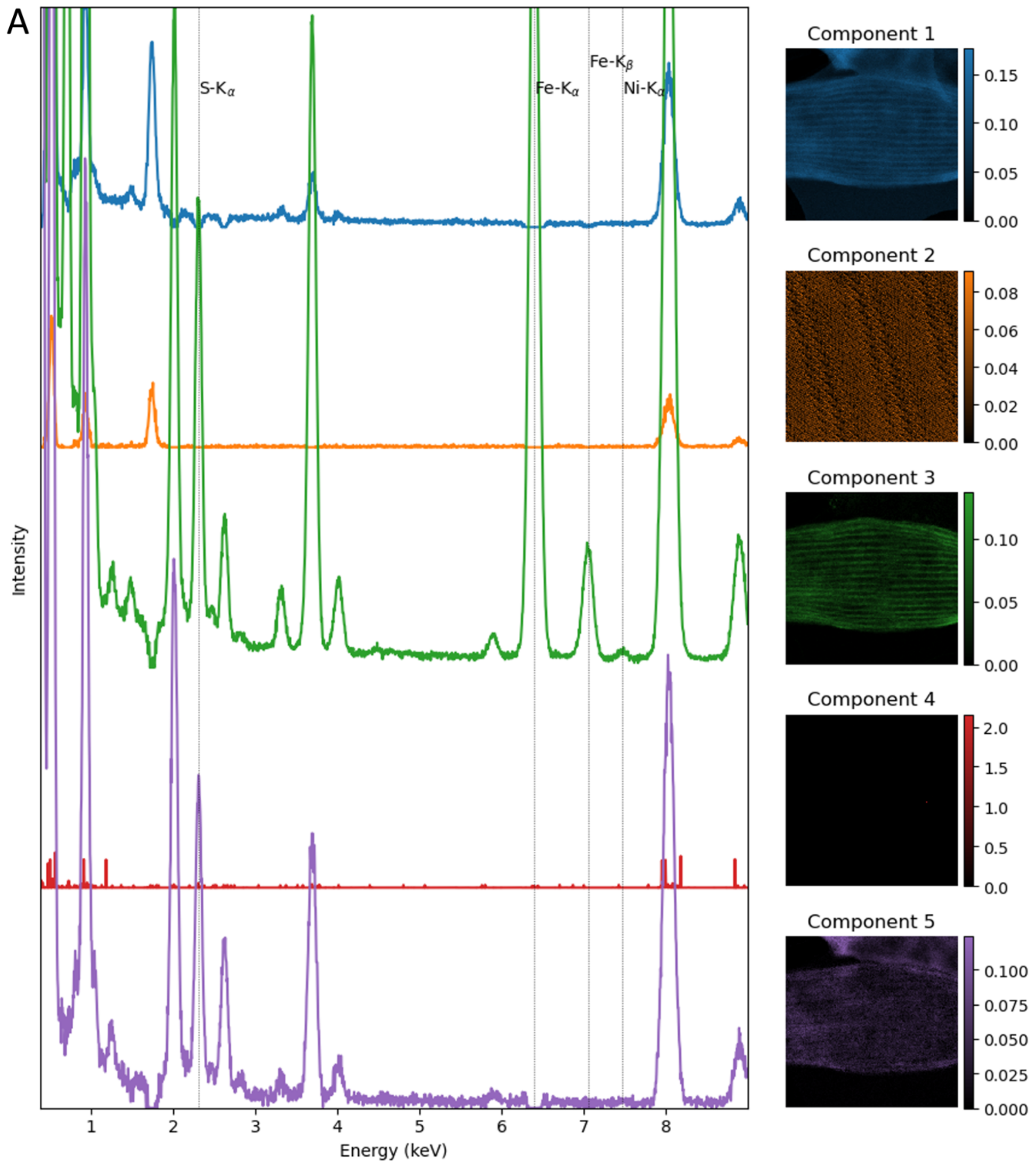
A
S-Kα
Fe-Kα
Fe-Kβ
Ni-Kα
Intensity
Energy (keV)
Component 1
Component 2
Component 3
Component 4
Component 5

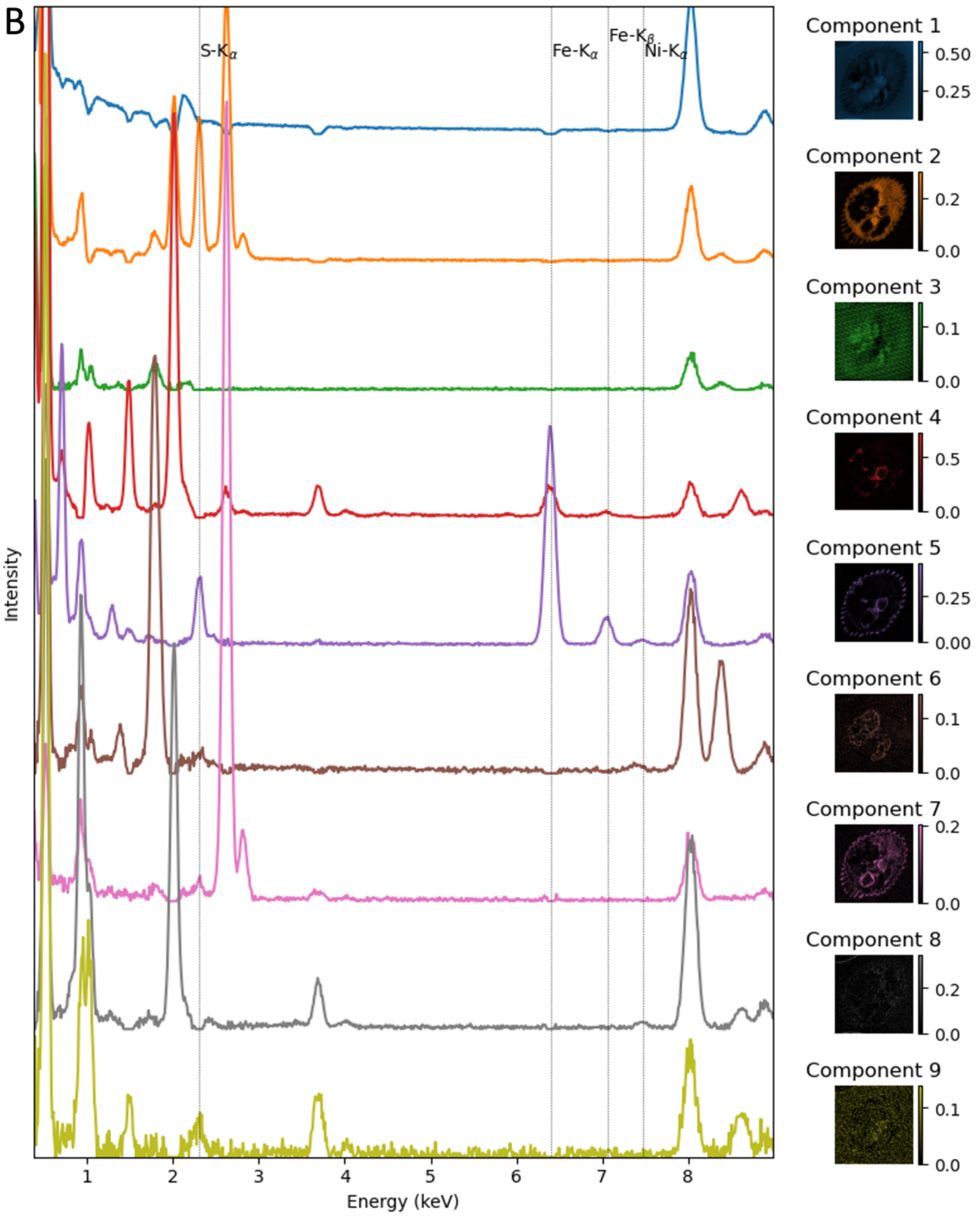
B
S-Kα
Fe-Kα
Fe-Kβ
Ni-Kα
Intensity
Energy (keV)
1 2 3 4 5 6 7 8
Component 1
0.50
0.25
Component 2
0.2
0.0
Component 3
0.1
0.0
Component 4
0.5
0.0
Component 5
0.25
0.00
Component 6
0.1
0.0
Component 7
0.2
0.0
Component 8
0.2
0.0
Component 9
0.1
0.0

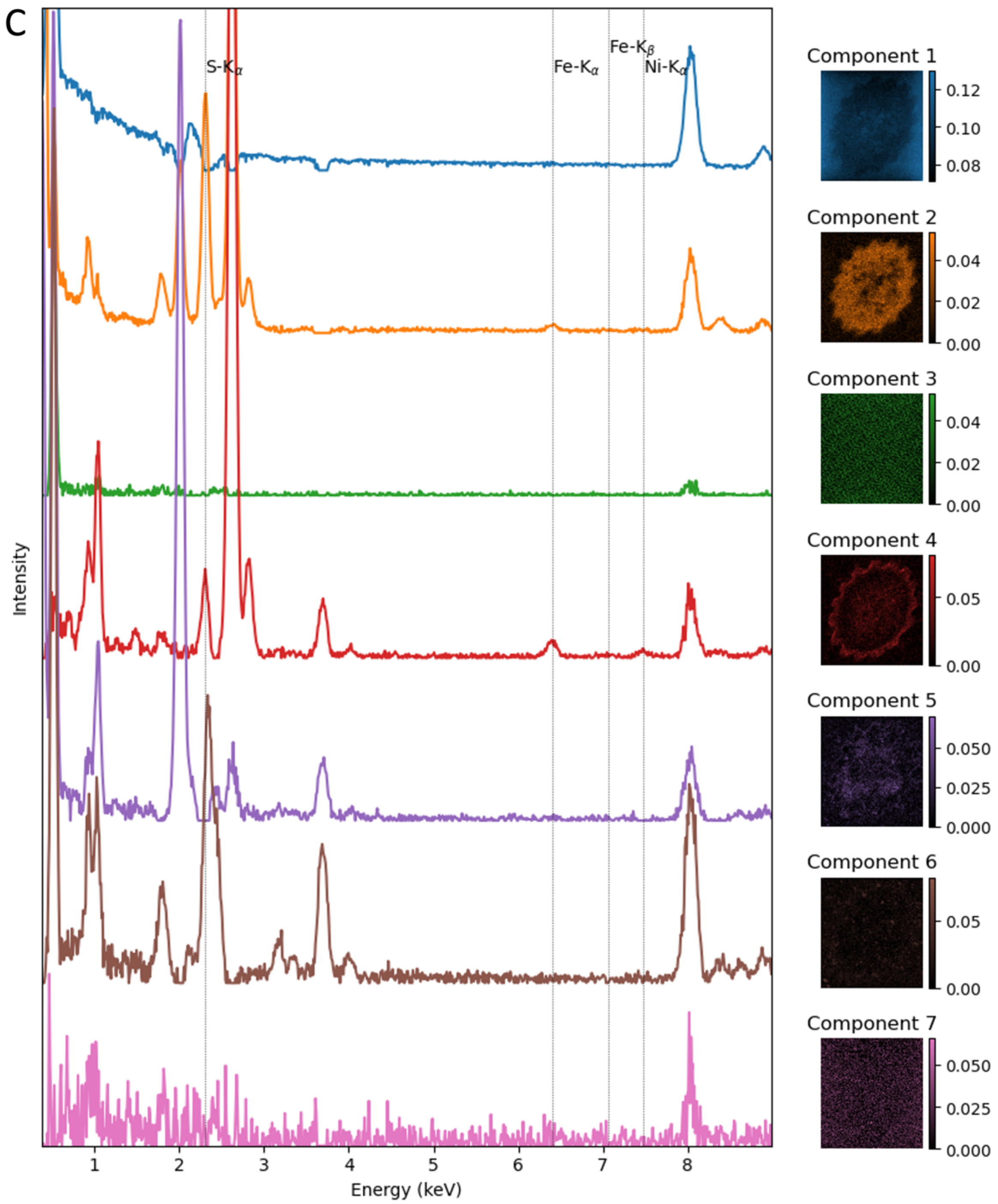


**Figure S4. A, B, C:** NMF decomposition of STEM-EDX spectrum images of an intact GS cell (**A**), an GS cross-section (**B**), and an RB cross-section (**C**). The extracted sum spectra for each component and relative intensities of components in the sample are shown on the left and right, respectively.

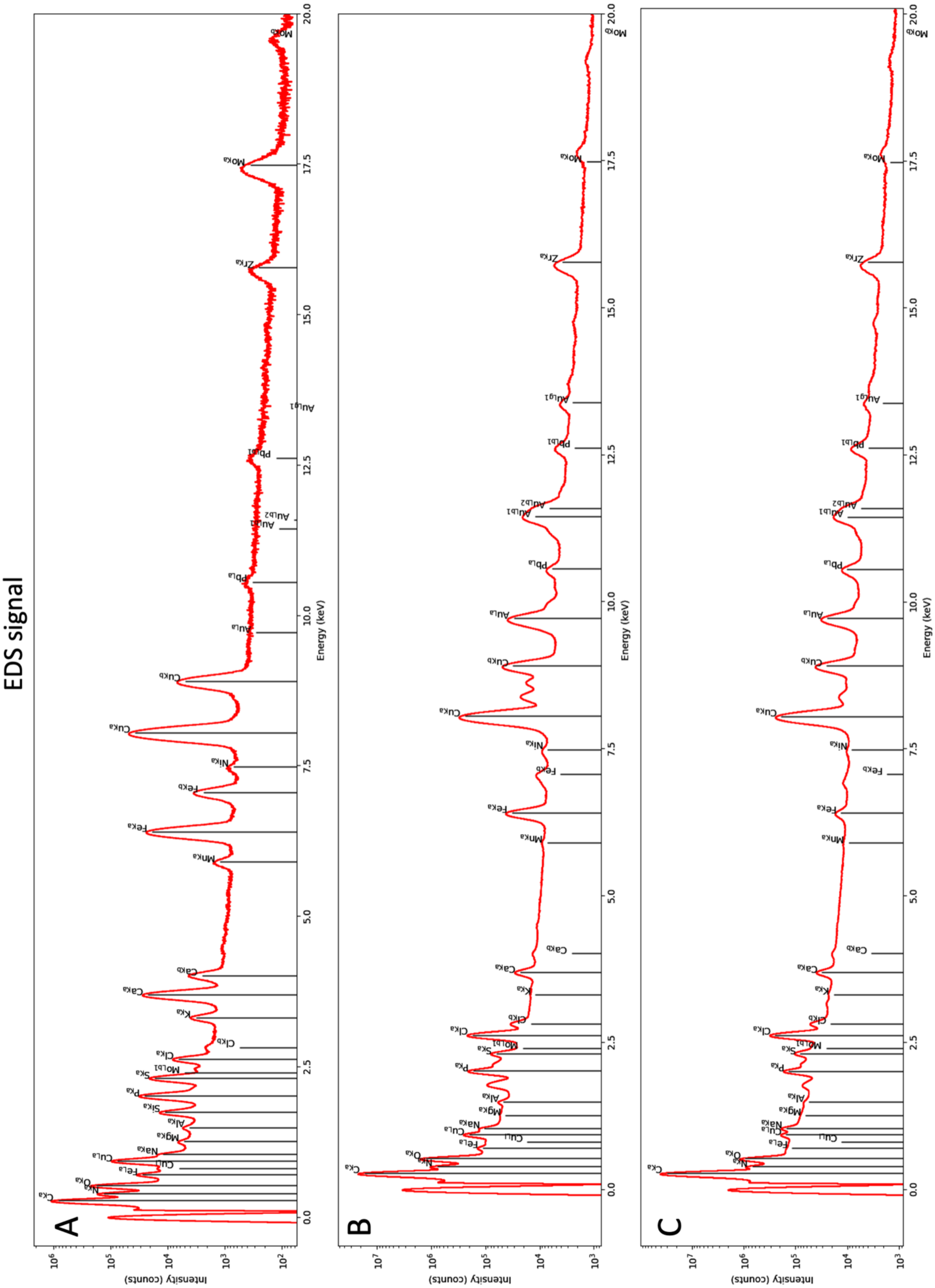


**Figure S5. A:** Summed spectrum of intact GS cell. **B:** Summed spectrum of GS cross-section. **C:** Summed spectrum of RB cross-section.

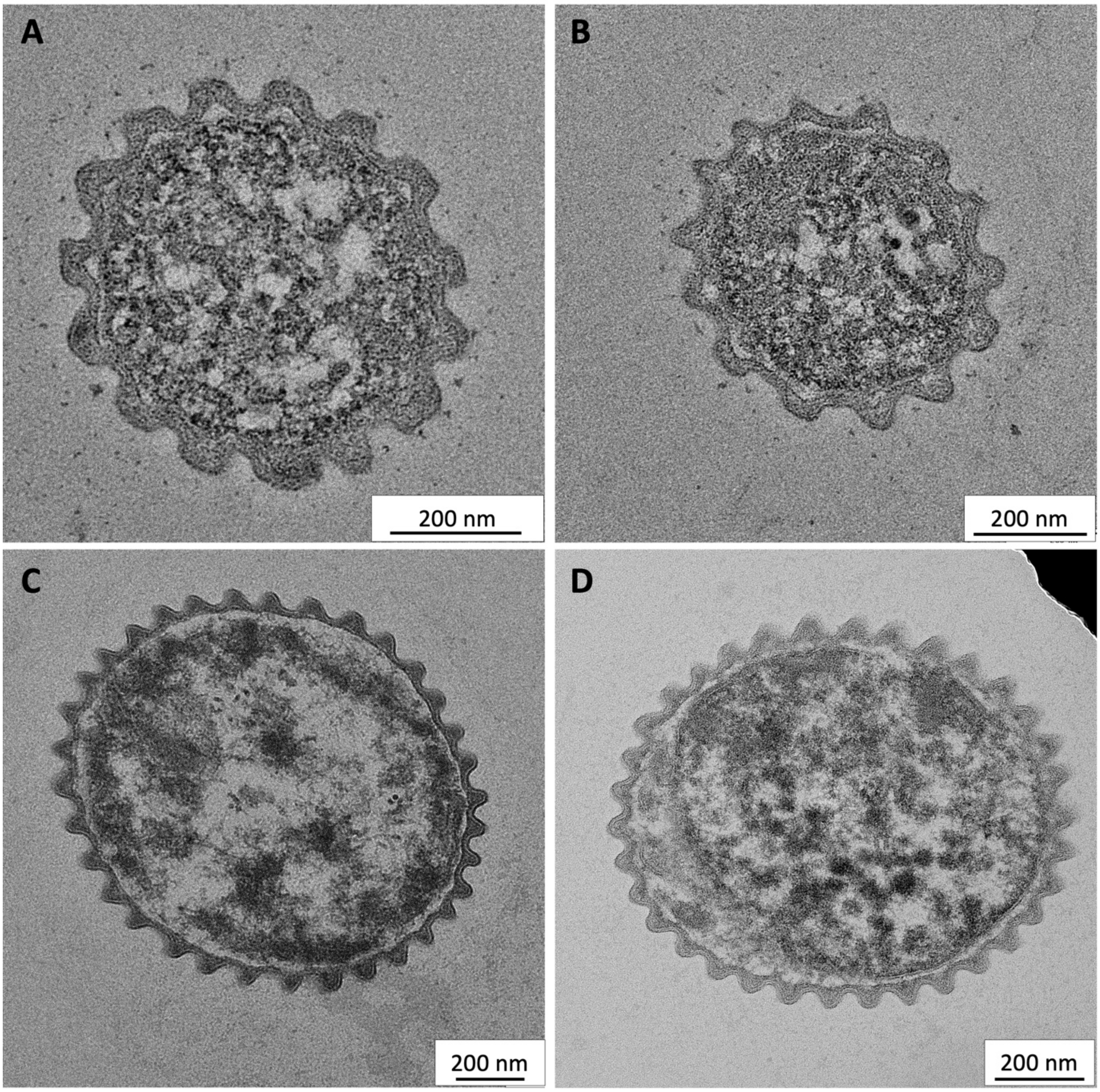


**Figure S6. TEM of plastic-embedded cross sections from cable bacteria filaments. A, B:** RB plastic cross-sections showing A: rectangular PCFs and B: both rectangular and rounded PCFs. **C, D:** GS plastic cross-sections with consistent rounded shape of the PCFs.

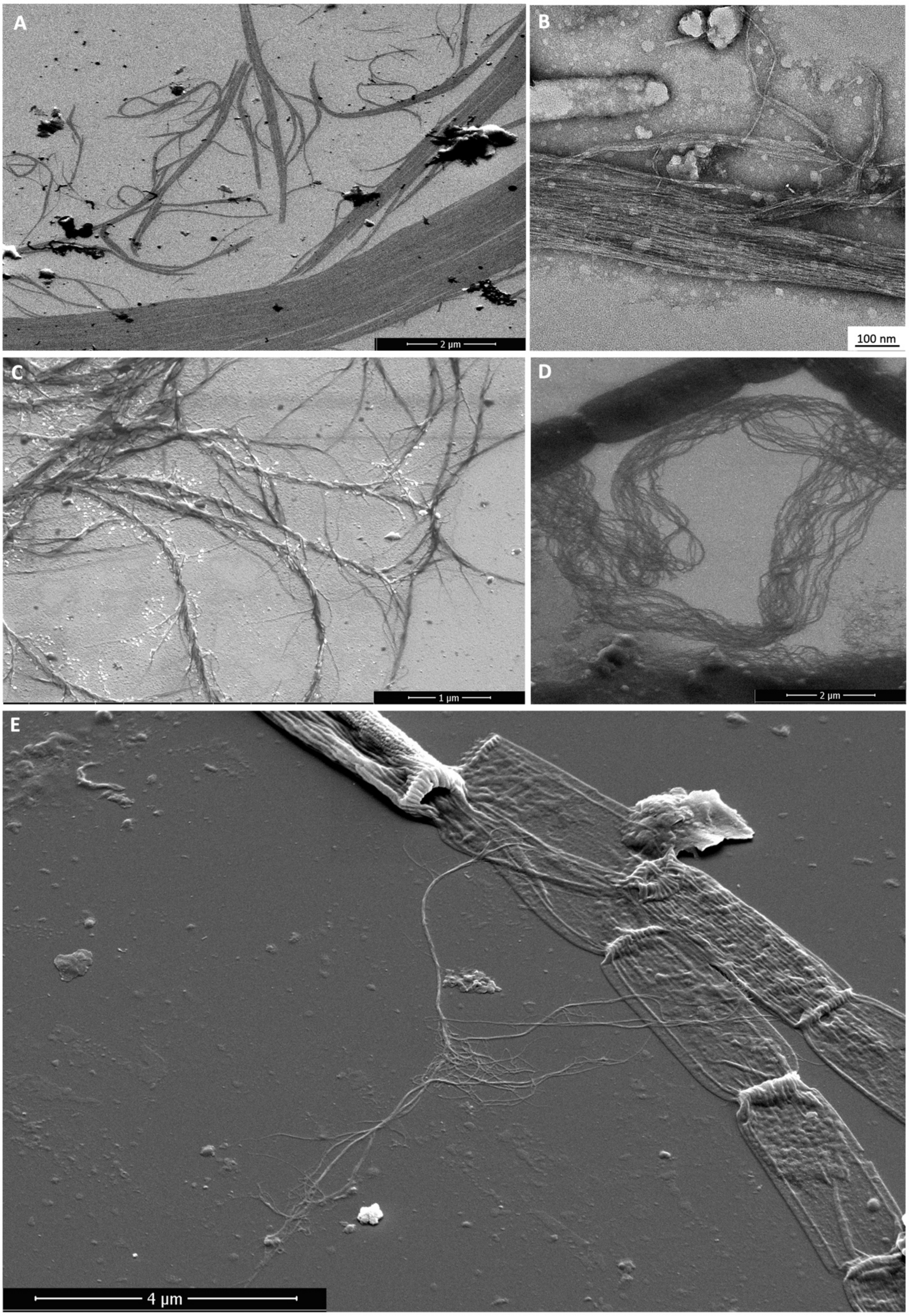


**Figure S7. SEM images of PCF strand components from different cable bacteria filaments. A, B:** Wild type marine cable bacteria released PSC after centrifugation at 12000g for 10min **C, E:** GS released PSC after harsh mechanical treatment **D:** Wild type marine cable bacteria from Hou beach released PSC after 100min incubation in SDS
